# ASTER: A Method to Predict Clinically Actionable Synthetic Lethal Genetic Interactions

**DOI:** 10.1101/2020.10.27.356717

**Authors:** Herty Liany, Anand Jeyasekharan, Vaibhav Rajan

## Abstract

A Synthetic Lethal (SL) interaction is a functional relationship between two genes or functional entities where the loss of either entity is viable but the loss of both is lethal. Such pairs can be used to develop targeted anticancer therapies with fewer side effects and reduced overtreatment. However, finding clinically actionable SL interactions remains challenging. Leveraging unified gene expression data of both disease-free and cancerous samples, we design a new technique based on statistical hypothesis testing, called ASTER, to identify SL pairs. We empirically find that the patterns of mutually exclusivity ASTER finds using genomic and transcriptomic data provides a strong signal of SL. For large-scale multiple hypothesis testing, we develop an extension called ASTER++ that can utilize additional input gene features within the hypothesis testing framework. Our extensive experiments demonstrate the efficacy of ASTER in identifying SL pairs with potential therapeutic benefits.

**CCS CONCEPTS:** **• Applied computing** → **Computational genomics**; **Health informatics**; • **Mathematics of computing** → **Hypothesis testing and confidence interval computation**.

**ACM Reference Format:** Herty Liany, Anand Jeyasekharan, and Vaibhav Rajan. 2021. ASTER: A Method to Predict Clinically Actionable Synthetic Lethal Genetic Interactions. In *Proceedings of ACM Conference*. ACM, New York, NY, USA, 10 pages. https://doi.org/10.1145/nnnnnnn.nnnnnnn

## 1 INTRODUCTION

Cancer is one of the leading causes of mortality and morbidity worldwide [40]. Cancer develops as a result of genetic alterations caused by endogenous and exogenous cellular processes; cancer cells gain selective advantage over healthy cells resulting in their ability to proliferate uncontrollably [23]. Many targeted therapies have been developed to interfere with specific molecular targets, e.g., driver genes, that are known to aid tumor growth and proliferation. However, cancer treatment continues to remain a challenge due to the heterogeneous nature of the proliferation that leads to substantial diversity in subtypes and treatment outcomes. The efficacy of targeted therapies has been found to be limited due to the emergence of acquired drug resistance and low prevalence of some alterations [10, 33, 60]. Further, the list of actionable alterations and suitable drugs to target known alterations, both remain incomplete [26, 38, 50, 52]. These difficulties have led to the investigation of alternative approaches to find drug targets.

Exploiting Synthetic Lethality is one such promising approach to discover effective therapeutic targets [39, 43]. A Synthetic Lethal (SL) genetic interaction is a functional relationship between two genes or functional entities where the loss of either entity is viable but the loss of both is lethal. Thus, the entities are likely to have redundant or compensatory roles that often give rise to signals of mutual exclusivity in genomic or transcriptomic data. The key idea used in targeted cancer therapeutics is that in a malignant cell, functionally disruptive mutation in one of the two genes (say, A) of an SL pair (A,B) leads to dependency on B for survival and cancer cells can be selectively killed by inhibiting B. Non-cancerous cells, that have A, survive even when B is inhibited. See Fig. 1 for a schematic. Thus targeted treatments based on Synthetic Lethality have the potential to be highly specific with fewer adverse effects, and may reduce overtreatment [39]. A well known SL pair is BRCA/PARP: mutations causing functional loss of BRCA1/2 genes leads to deficiency of DNA Damage Response mechanism and dependence on the protein PARP1/2 [11]. Drugs based on PARP inhibitors have been found to be effective for breast and ovarian cancers [1, 49].

**Figure 1:**
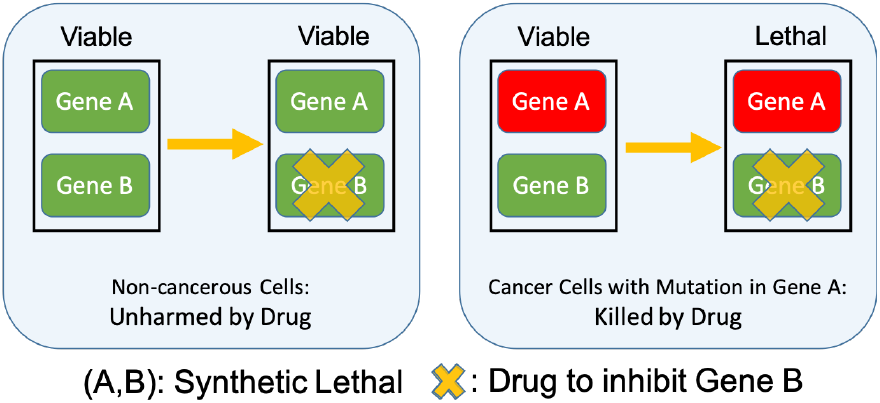
Synthetic Lethality: gene A or gene B can ensure cell survival, but loss of both is lethal.

Despite the concept being decades old (from genetic studies in fruit fly and yeast, e.g., [5, 35]), SL interactions in humans remain largely unknown. In fact, PARP-inhibitors are the only approved drugs based on SL [10]. Genome-wide screens have been developed, e.g., RNA interference screens and CRISPR screens, to identify potential SL pairs but they are costly and labour-intensive. There are several challenges in the identification of SL interactions: first, since these genetic interactions are lethal, mutant recovery and identification become difficult; second, many SL pairs are conditionally dependent and may not be conserved in all genetic backgrounds or in different cellular conditions and third, large number of gene pairs need to be queried to identify SL interactions [39]. Hence, there is a need to develop computational methods to identify and prioritize potential SL pairs, as candidates that can be functionally analyzed through genome-wide screens.

Many computational methods have been developed for SL prediction. Broadly, they can be divided into machine learning and statistical methods. Machine learning methods face the challenge of scarce labels since very few human SL pairs are experimentally confirmed. Statistical approaches, such as DAISY [28] and ISLE [31], have the advantage of not relying on labelled data. Moreover, they are based on well-defined biological hypotheses and are easily interpretable. Both DAISY and ISLE procedures consist of multiple hypothesis tests, that use multiple datasets. For instance, DAISY uses pan-cancer Somatic Copy Number Alteration (SCNA) profiles, gene expression data and mutation profiles (from The Cancer Genome Atlas (TCGA) [54]) in addition to shRNA-based functional essentiality profiles. The aim of their tests is to discover underlying patterns in the data from which the mutual exclusivity pattern indicative of SL can be deduced. More details are in section 2. We postulate that the use of disease-free samples in mining underlying patterns is important to identify clinically actionable SL pairs. This is because both the genes in the SL pair have to be functional in non-cancerous cells, as shown in Fig. 1, for the target drug to be effective. To our knowledge, the use of disease-free samples in finding SL pairs has not been evaluated in any previous method.

RNA-Seq expression data from disease-free tissues are available in the Genotype Tissue Expression (GTEx) project for over 53 tissues [34]. Data in GTex has been unified with cancer tissues from TCGA, after re-alignment, re-quantification of gene expression and successful correction for study-specific biases [20, 53]. Thus, GTEx provides tissue-specific reference expression levels for comparison with the expression levels found in cancer. In this paper, we leverage GTEx data to design a new technique called ASTER (**A**nalysis of **S**ynthetic lethality by comparison with **T**issue-specific disease-free g**E**nomic and t**R**anscriptomic data) to identify potential SL gene pairs. ASTER is based on statistical hypothesis testing, and the tests in ASTER follow directly from the mutual exclusivity pattern in the definition of Synthetic Lethality. Using only SCNA and RNA-Seq data from GTEx and TCGA, and a straightforward formulation, ASTER is a relatively simple method that outperforms previous methods based on hypothesis testing, in our experiments.

For large-scale multiple hypothesis testing, we also develop an extension called ASTER++ that uses AdaFDR [58] to adaptively find a decision threshold based on additional gene features. Similar to machine learning based methods, ASTER++ can utilize gene-specific features in its predictions, but without their limitation of requiring labelled data to learn from. Moreover, it retains the interpretability of statistical hypothesis testing, while leveraging on AdaFDR’s scalability and flexibility. Our experiments demonstrate the efficacy of ASTER and ASTER++ in accurately identifying SL pairs that are therapeutically actionable in multiple cancers.

## 2 RELATED WORK

Previous methods to identify SL pairs can be broadly categorized into machine learning and statistical methods. Machine learning methods, including those based on network analytics, rely on labelled data to predict SL pairs, e.g., [7, 12, 27, 32]. However, these methods face the challenge of scarce labels due to lack of experimental confirmation. Some approaches have developed models that can incorporate information from experimentally confirmed yeast SL pairs [15, 46, 55]. These methods face several hurdles due to non-overlapping genes and/or partially overlapping functions of genes. Several models use binary classifiers trained on data where the evidence of negative labels, i.e., those pairs that are *not* SL, are not confirmed through screens. In general, such positive-unlabelled learning tasks, with scarce positive labels can be challenging [4].

Statistical approaches, that do not rely on labelled data, such as DAISY [28] and ISLE [31], are popular alternatives based on well-defined biological hypotheses. For a gene pair, (A,B), DAISY applies three statistical inference procedures. The first test, uses multiple SCNA and gene expression datasets to detect gene pairs that are infrequently co-inactivated. A Wilcoxon rank sum test is used to check if B has significantly higher SCNA level in samples where A is inactive, compared to remaining samples in each dataset. The input pair passes the first test if it passes the rank sum test in any of the datasets. The second test uses multiple shRNA essentiality datasets, SCNA and gene expression profiles, to identify pairs where underexpression and low copy number of a gene induces essentiality of the partner gene. A Wilcoxon rank sum test is conducted to test if B has significantly lower essentiality (lower score implies more essential), in samples where A is inactive compared to rest of the samples in each dataset. The input pair passes the second test if it passes the rank sum test in any of the datasets. The third test checks for significant co-expression in transcriptomic datasets, with the assumption that SL pairs, participating in related biological processes, are likely to be coexpressed. Positive Spearman’s correlation coefficient in at least one of the datasets is required for a pair to pass the third test. The input pair is considered SL if all three tests are passed.

ISLE is designed with a different aim – to obtain clinically relevant SL pairs from an initial (larger) collection of potential SL pairs. They also apply three statistical procedures, but unlike DAISY, the tests are done in a sequential manner. In the first test, gene expression and SCNA data is used to identify candidate gene pairs with significantly infrequent co-inactivations, signifying under-represented negative selection. This is found using a hypergeometric test on gene expression and SCNA data from tumor samples. Second, from the selected gene pairs in the first test, a gene pair is selected if its co-inactivation leads to better predicted patient survival, compared to when it is not co-inactivated. Survival probability is predicted using Cox proportional hazard model while controlling for confounding factors such as cancer type, patient age, gender, ethnicity and an index based on SCNA data. In the final step, from the selected gene pairs in the second test, only pairs with high phylogenetic similarity are retained, assuming co-evolution of functionally interacting genes. Thus, apart from SCNA and gene expression profiles, ISLE also uses clinical data and phylogeny information.

In comparison, the method we develop, ASTER, utilizes fewer datasets and has a simpler hypothesis testing framework. ASTER directly mines mutual exclusivity patterns that have been shown to provide strong signal for SL in previous studies, e.g. [45]. The tests in ASTER use only SCNA and gene expression data from cancerous and disease-free tissues, from TCGA and GTEx respectively. The tests in ASTER are also tissue-specific that is possible with availability of tissue-specific gene expression in GTEx. Thus, it is in alignment with the differences observed in the molecular and phenotypic characteristics, in particular genomic alterations that promote tumorigenesis, across cancers of different tissues [22, 42].

## 3 OUR METHOD: ASTER

ASTER (**A**nalysis of **S**ynthetic lethality by comparison with **T**issue-specific disease-free g**E**nomic and t**R**anscriptomic data), comprises sequential application of three tests for a candidate gene pair (A,B). The main idea is to check if there are tissue-specific cancer samples where (A,B) exhibits a mutual exclusivity pattern, i.e., where A is significantly up-regulated and simultaneously B is significantly down-regulated and the gene expressions are significantly different. Using disease-free tissues as reference, we find samples where A is significantly up-regulated (Test T1) and where B is significantly down-regulated (Test T2). Finally, we test if the expression levels are significantly different (Test T3). During sample selection we also consider SCNA data to account for copy number amplifications and deletions. We now give a formal description of the tests.

Let *S* _(*A* ↑)_ be the set of tissue-specific samples (from TCGA) with high copy number (SCNA *>* 1) for gene A. Let *S* _(*B* ↓∈ *A* ↑)_ ⊂*S*_(*A* ↑)_ be the set of samples in *S*_(*A*↑)_ with low copy number (SCNA *<* 1) for gene B. Let *N* denote non-cancerous samples of the same tissue type (from GTEx). The following tests are applied sequentially:

T1: We test if the expression levels of gene A in *S* _(*A* ↑)_ is significantly higher than the expression levels of A in *N*.

T2: We then test if the expression level of gene B in *S*_(*B* ↓∈ *A* ↑)_ is significantly lower than the expression levels of B in *N*.

T3: Finally, we test if the expression levels of gene A in *S* _(*A* ↑)_ is significantly higher than the expression levels of gene B in *S* _(*B* ↓∈*A*↑)_.

Figure 2 shows a schematic of the three tests in ASTER. We use the non-parametric Wilcoxon rank sum test for each of the three tests. Fisher’s method [17] is used to obtain a single p-value by combining the p-values from the three independent tests. Note that due to the sequential manner of testing, the application of ASTER on gene pairs (A,B) and (B,A) may yield different results. ASTER explicitly tests for up-regulation and amplification in the first gene and simultaneous down-regulation and deletion in the second gene. Further, the order of tests T1 and T2 can be reversed, i.e., we can test for down-regulation in the first gene using tissue-specific cancer samples with low copy number *S* _(*A* ↑)_ and then test for up-regulation in the second gene using samples *S* _(*B* ↓∈ *A* ↑)_that have high copy number in B among the samples *S* _(*A* ↑)_. The third test remains the same. Thus, for an input pair (A,B), there are four possible ways of testing depending on the sequence of the genes and the sequence of the first two tests. We denote these four cases by: (*A* ↑ *B* ↓), (*B* ↑ *A* ↓), (*A* ↑ *B* ↓), (*B* ↑ *A* ↓). Unless mentioned otherwise, we test all four cases and an input pair is predicted to be SL if it passes all three tests in at least one of the four cases.

**Figure 2:**
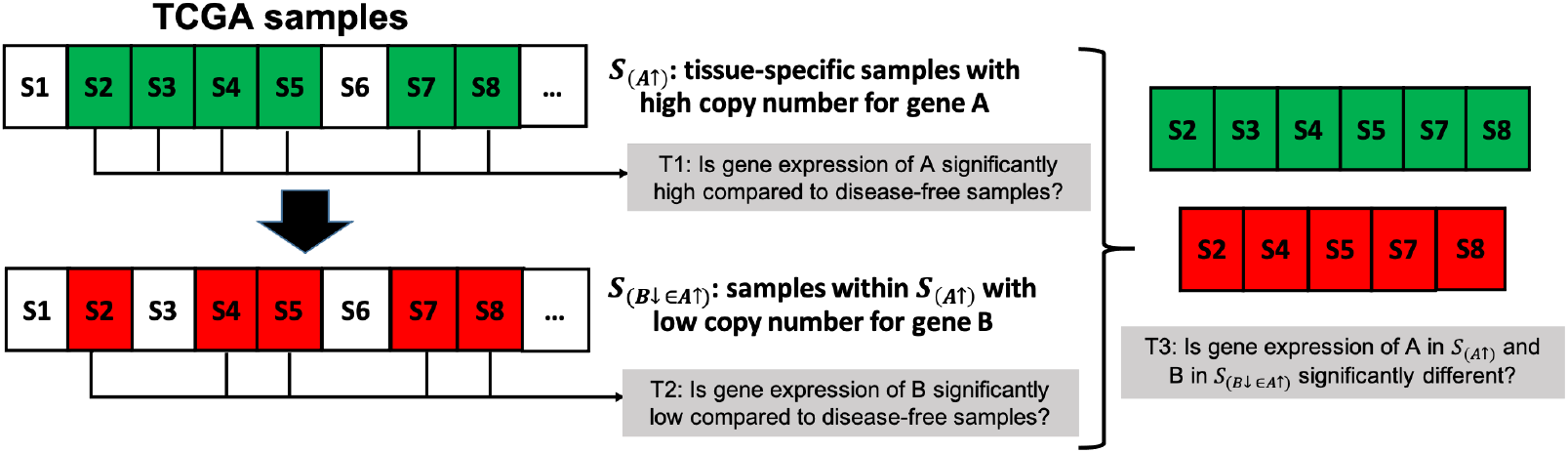
Overview of ASTER. T1: Green color indicates samples in *S*_(*A*↑)_ where gene A is significantly up-regulated (compared to disease-free GTex samples). T2: Red color indicates samples in *S*_(*B* ↓∈ *A* ↑)_ where gene B is significantly down-regulated (compared to disease-free GTex samples). T3: Gene expression values of selected samples are compared to conduct test T3.

**Figure 3:**
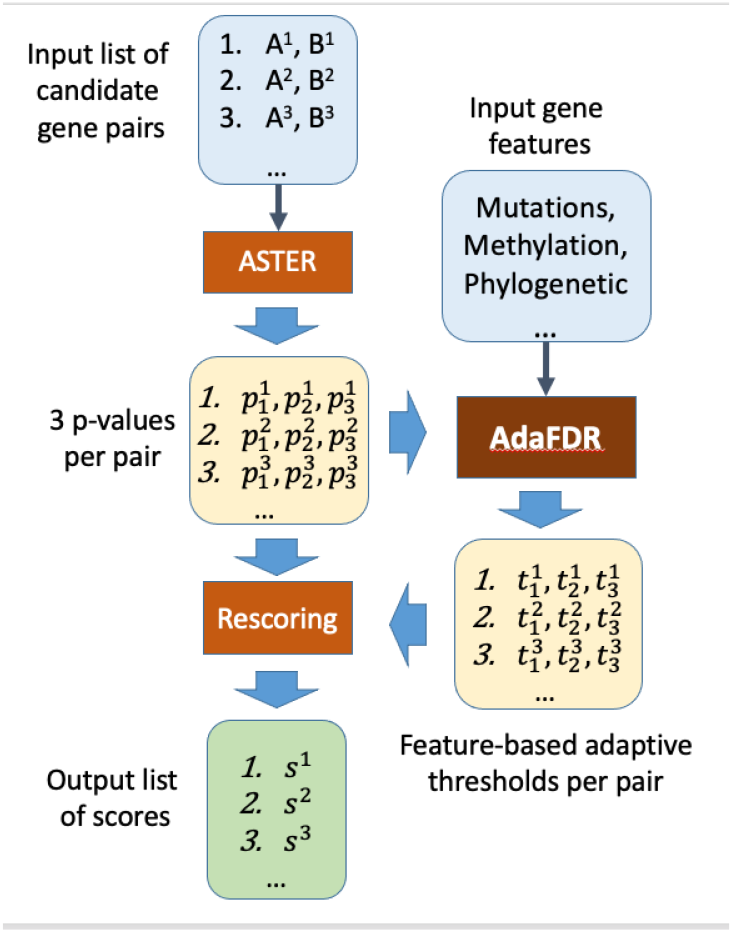
ASTER++ pipeline for large-scale multiple testing and use of additional features of input gene pairs.

We do not use mutation-based data in ASTER. Due to intra- and inter-tumor heterogeneity of cancer samples, mutations in cancer samples may be hard to validate and prone to high error rate, with higher false-positive rates for driver genes [2]. Empirically we find that gene expression and SCNA profiles, especially through comparison with disease-free expression levels, provide sufficiently robust signal for SL detection.

### 3.1 ASTER++

To enable large-scale multiple hypothesis testing and the use of additional known covariates about the gene pairs, we combine ASTER with AdaFDR [58]. AdaFDR adaptively finds a decision threshold from covariates at a user-specified False Discovery Proportion. For a candidate list of gene pairs, their features and corresponding p-values (obtained from ASTER in our case), AdaFDR learns a decision threshold that depends on the covariates. Thus, instead of a fixed threshold used in previous hypothesis testing based methods, through the use of AdaFDR, we can obtain a covariate-dependent threshold that may be different for each gene pair. While the adaptive threshold can be directly used to predict SL for a specific gene pair, it cannot be used to rank multiple input gene pairs. We address this through the following re-scoring strategy.

Let 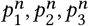 be the p-values from the three tests of ASTER for the *n*^th^ gene pair. With a fixed threshold (e.g., *t* = 0. 01) we can choose all the pairs with 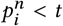 for *i* = 1, 2, 3 as the predicted SL pairs. They can be ranked using the single p-value obtained after applying Fisher’s method. When an adaptive covariate-dependent threshold is learnt from AdaFDR, the *n*^th^ gene pair has three different thresholds 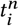, one for each p-value 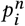 for *i* = 1, 2, 3. We can consider 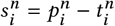 to be a score which indicates how far each p-value is from its own threshold, with a lower value indicating higher significance. For the *n*^th^ gene pair and the *i*^th^ test there is a decision value 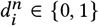 that is set to 1 if 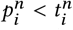. Only those gene pairs are selected that pass all three tests, i. e., when 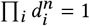. The selected gene pairs can be ranked using the score 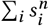 with a lower value indicating higher probability of being SL.

We call this combined method of using ASTER, AdaFDR and rescoring as described above ASTER++. Similar to machine learning based methods, ASTER++ can utilize gene-specific features in its predictions, without their limitation of requiring labelled data to learn from. Moreover, it retains the interpretability of ASTER’s statistical hypothesis testing, while leveraging on AdaFDR’s scalability and flexibility of large-scale multiple hypothesis testing.

## 4 EXPERIMENTS

We first compare the performance of ASTER with that of DAISY and ISLE, on identifying SL pairs in benchmark datasets and on drug efficacy prediction tasks that indirectly evaluate SL in section 4.1. We then evaluate the predicted pairs from a specific dataset qualitatively in section 4.2.

### 4.1 Performance Comparison on SL Prediction

In this section, we compare the performance of ASTER and ASTER++ with two state-of-the-art methods, DAISY and ISLE, that are based on hypothesis testing. We conduct three experiments. First, we compare the methods on benchmark datasets where SL interactions have been validated using CRISPR and/or shRNA screens. In the next two experiments, we utilize drug response screens on model cell lines to evaluate the predictions through the observed inhibitory effects of the drugs on cell lines with loss of function alterations in genes from the predicted SL pairs.

#### 4.1.1 Parameter Settings for Methods Used

We describe here the settings used for ASTER, ASTER++, DAISY and ISLE in our experiments.

For the three tests in DAISY, we follow the procedure described by the authors in [28]. We obtained SCNA, mRNA gene expression data and mutation profiles from multiple cancers (BRCA, LUAD, CESC, KIRC, KIRP, KICH, AML) in TCGA [54], using cBioPortal [18] and Firehose. Essentiality profiles are based on those curated in [37]. DAISY uses a p-value cutoff of 0.05 after Bonferroni correction for multiple hypotheses testing. For ISLE, we use the software and data provided by them, using related cancer type data. We obtained phylogenetic similarity for 86 species using the phylogenetic profile database [47]. ISLE uses FDR *<* 0.2 based on Benjamini-Hochberg and a cut-off of 0.5 to determine phylogenetically linked pairs. ASTER, DAISY and ISLE, each yield three p-values per gene pair, that are combined using Fisher’s method [17].

As mentioned in section 3, for each input gene pair, 4 cases are considered. A pair is predicted as positive if any of the 4 cases passes the test of ASTER, i.e., obtains all 3 p-values less than 0.01. A pair is predicted negative if all 4 cases do not pass the 3 tests of ASTER. For ASTER++, the covariates used are the following for each gene in the input pair: (1) loss of function mutation counts, (2) phylogenetic score, (3) methylation (HM450) beta-values and (4) the number of samples that exhibit up-/down-regulation for each gene based on the output results from ASTER. Loss of function mutation counts is the sum of non-synonymous mutations in each gene (excluding synonymous mutations such as ‘Silent’, ‘Intron’, ‘3’UTR’, ‘5’UTR’, ‘IGR’, ‘lincRNA’ and ‘RNA’). Methylation value refers to human’s gene methylation (HM450) beta-values (aberrant DNA methylation, i.e., hyper- or hypo- methylation, has been implicated in many disease processes, including cancer). Both loss of function mutations and methylation value were retrieved from cBioportal [18]. The phylogenetic score of a gene describes the relative sequence conservation or divergence of orthologous proteins across a set of reference genomes and has been used in several tasks such as gene annotation and function prediction. The phylogenetic score was retrieved from phylogenetic profile database by [47].

#### 4.1.2 Predictive Accuracy on Benchmark Datasets

We use 2 benchmark datasets wherein SL interactions have been validated using CRISPR and/or shRNA screens. The first a benchmark dataset containing breast cancer SL pairs from SynLethDB [21] where we select those pairs that have evidence of SL from multiple sources including text mining, genomeRNAi and shRNA screens. The second dataset has 15,313 SL pairs from the functional study performed by [24] on leukemia cell lines. To form a negative set, we follow the procedure in [32]. We randomly select genes from the HGNC database [9] after excluding genes reported in any SL interaction in SynLethDB and those reported to be essential in [37, 51]. We use 1,000 and 15,313 gene pairs as negative samples. in the datasets respectively.

We report the counts of True Positives (TP), True Negatives (TN), False Positives (FP) and False Negatives (FN). Each method comprises three tests and we show the counts for each test separately. The final counts are obtained by considering those pairs that pass all three tests in DAISY, ASTER and ASTER++. In the case of ISLE, we follow their sequential filtering approach to obtain the final counts: only those pairs that pass the first test are considered for the second test, and only those pairs that pass the second test are considered for the third test, and the final counts show the results after the third test.

While there is evidence of SL through experimental screens like CRISPR for the positive pairs in our datasets, for the randomly selected negative pairs in our data there is no conclusive evidence of their *not* being SL. Hence, they should be considered untested or unlabelled. This setting is similar to many other applications that have positive-unlabelled data, such as recommendation systems. In such cases, metrics that use True Negatives or False Positives, e.g., Precision or AUC, are not reliable. A standard approach is to evaluate predictive algorithms through Recall@N, which is defined as the proportion of pairs correctly identified as True Positives, among the total list of SL pairs (245 in the first dataset and 15,313 in the second dataset), in a ranked list containing the best scoring *N* items. In our experiments, the p-values are used as scores, with lower values indicating better scores.

#### Results

Table 1 shows the final counts for each method on the breast cancer dataset. There are 3 tests in each method, and the counts for each of these are also shown. As expected, ISLE is the most stringent (by design) and the second test does not pass any of the positive pairs. In the final row, we show all the true negatives and false negatives from both the first two tests for ISLE. Very few pairs pass the 3 tests of DAISY and ASTER – only 3 for DAISY and 5 for ASTER. ASTER++ can detect many more pairs but also predicts many false positives.

**Table 1:** Breast Cancer Data: TP, FP, TN and FN counts within each test and the final results.

|  | DAISY |  |  |  | ISLE |  |  |  | ASTER |  |  |  | ASTER++ |  |  |  |
| --- | --- | --- | --- | --- | --- | --- | --- | --- | --- | --- | --- | --- | --- | --- | --- | --- |
|  | TP | FP | TN | FN | TP | FP | TN | FN | TP | FP | TN | FN | TP | FP | TN | FN |
| 1st | 222 | 721 | 279 | 23 | 231 | 800 | 200 | 14 | 241 | 777 | 223 | 4 | 244 | 796 | 204 | 1 |
| 2nd | 27 | 46 | 954 | 218 | 0 | 0 | 1000 | 245 | 11 | 4 | 996 | 234 | 78 | 77 | 923 | 167 |
| 3rd | 37 | 53 | 947 | 208 | 139 | 483 | 517 | 106 | 9 | 8 | 992 | 236 | 101 | 133 | 867 | 144 |
| final | 3 | 3 | 997 | 242 | 0 | 0 | 1000 | 245 | 5 | 1 | 999 | 240 | 61 | 31 | 969 | 184 |

Fig. 4 shows the performance in terms of Recall@N of ASTER, DAISY, ISLE and ASTER++. ASTER outperforms both DAISY and ISLE at values of N up to 120 and is comparable at values of 150 and higher. When covariates are available, ASTER++ boosts the performance of ASTER particularly at higher values of N. Table 2 shows the true positive counts for each method at each value of N. Comparing this result with Table 1 shows that for a fixed p-value cutoff DAISY, ASTER and ISLE pass very few of the true pairs. However, if we consider the top-scoring pairs from each method, without considering a fixed cutoff value, then all the methods find some true SL pairs in their respective ranked lists. Thus, among the top 30 predicted pairs in their respective lists, ISLE finds 10 true SL pairs and DAISY finds 17. ASTER outperforms both of them and finds 22 true SL pairs in its top 30 while the use of additional features leads to ASTER++ finding 27 true pairs in its top 30. As we go down the ranked list, the number of additional SL pairs found reduces for all methods.

**Table 2:**
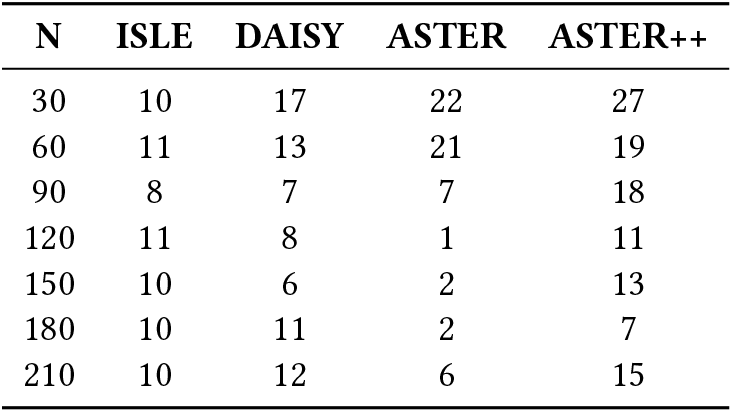
TP counts in Top N highest scoring predicted pairs for each method; used for Recall@N (Fig. 4). Each line after the first shows the number of additional pairs identified.

| N | ISLE | DAISY | ASTER | ASTER++ |
| --- | --- | --- | --- | --- |
| 30 | 10 | 17 | 22 | 27 |
| 60 | 11 | 13 | 21 | 19 |
| 90 | 8 | 7 | 7 | 18 |
| 120 | 11 | 8 | 1 | 11 |
| 150 | 10 | 6 | 2 | 13 |
| 180 | 10 | 11 | 2 | 7 |
| 210 | 10 | 12 | 6 | 15 |

**Figure 4:**
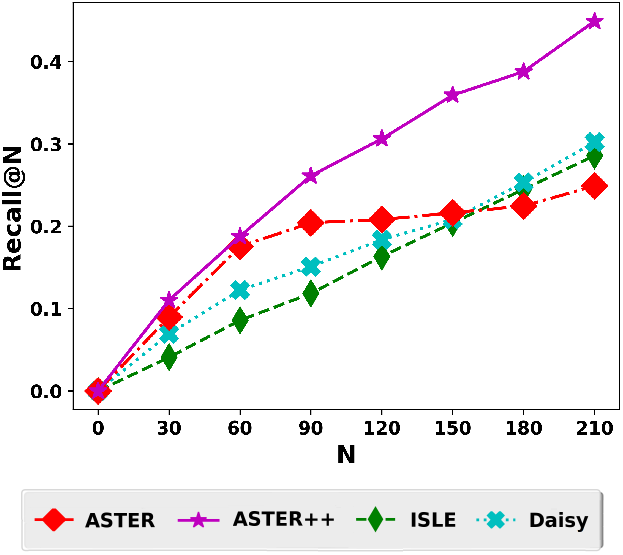
Recall@N of ASTER, ASTER++, ISLE and Daisy in 245 SL pairs in breast cancer from SynlethDB.

On the second leukamia dataset, the final counts from the methods are very similar to those in the SynlethDB dataset: most of the input pairs do not pass the tests of DAISY, ISLE or ASTER. One important difference we observed was that most p-values were not only insignificant but also close to 1. The counts are shown in Table 9 in the Appendix. The use of AdaFDR did not result in much difference in the adaptive threshold. Thus, ASTER++ results were nearly the same as those from ASTER. This may be due to the high and similar p-values. The performance with respect to Recall@N is more illustrative and shown in Fig. 5 with the TP counts in Table 3. The curve for ASTER++ coincides with that of ASTER and is not visible. We observe that ASTER (and ASTER++ that had the same results) consistently predicts true SL pairs in the first 240 pairs ranked by the p-values. The performance of DAISY is also good but not better than that of ASTER, while ISLE finds the least number of true pairs in its ranked list.

**Table 3:** TP counts in Top N highest scoring predicted pairs for each method; used for Recall@N (Fig. 5). Each line after the first shows the number of additional pairs identified.

| N | ISLE | DAISY | ASTER/ASTER++ |
| --- | --- | --- | --- |
| 30 | 24 | 26 | 30 |
| 60 | 20 | 26 | 30 |
| 90 | 20 | 26 | 30 |
| 120 | 16 | 21 | 30 |
| 150 | 19 | 24 | 30 |
| 180 | 19 | 24 | 30 |
| 210 | 17 | 22 | 30 |
| 240 | 18 | 25 | 30 |

**Figure 5:**
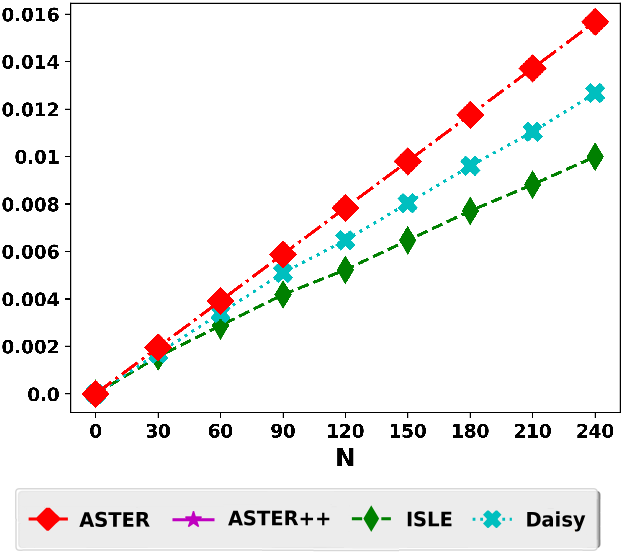
Recall@N of ASTER, ASTER++, ISLE and Daisy in 15,313 SL pairs from leukemia cell lines.

#### 4.1.3 Clinical Relevance of Predicted SL Pairs

In this experiment, we consider 26 target genes in the in-vitro drug response screen of 24 drugs on 500 cancer cell lines from the Cancer Cell Line Encyclopedia (CCLE) downloaded from [20]. These cell lines overlap with the cell lines in the Genomics of Drug Sensitivity in Cancer (GDSC) drug response database [57] that we use for our analysis. Among these cell lines, 30 are from Breast Cancer and 19 are from Stomach Cancer. We combine these 26 target genes with 32,018 genes from the human genome to form a total of 832,468 candidate (target-partner) gene pairs. These candidate pairs are used as input to ASTER, DAISY and ISLE. We do not use ASTER++ in this experiment since we want to specifically evaluate the mutual exclusivity pattern mined through ASTER’s tests. In this experiment, we use the same p-value cutoff value of 0.05 for all tests in all 3 methods. The predicted SL pairs are used for subsequent evaluation as follows.

GDSC provides results from their statistical analysis that correlates drug sensitivity with various genomic features such as mutations, amplifications and deletions of common cancer genes. ANOVA is used to identify important features associated with drug response across all cell lines. For each drug, a feature list is provided comprising genomic features, transcripts and tissue with an effect size assigned to each. Features with negative effect size are associated with drug sensitivity and features with positive effect size are associated with drug resistance. We use the associations provided between drug effect and the low copy number feature of genes in our analysis.

The principle behind the evaluation is from the concept of SL. We consider predicted SL pairs (A,B) where gene A is from the list of 26 target genes and gene B is from the list of 32,018 partner genes. If (A,B) is a SL pair and the partner gene B is associated with functional loss, then targeting gene A would be effective in inhibiting cancer growth. Thus, to evaluate the clinical relevance of the predicted SL pairs, we check the number of partner genes that are associated with loss of function through the low copy number feature in GDSC. Among those that are associated with low copy number, we find the proportion that are strongly associated with drug sensitivity (through negative effect size). This proportion provides indirect evidence of the predicted pairs being SL.

#### Results

Table 4 shows the number of pairs that passed all three tests of ASTER and DAISY: 3040 and 2766 pairs in Breast Cancer (BRCA); 1278 and 1228 pairs in Stomach Cancer (STCA). None of the candidate pairs pass all three tests of ISLE. These predicted pairs are used for subsequent evaluation. Table 4 also shows that very few predicted pairs have partners that are associated with loss-of-function (LoF) feature in GDSC – 47 out of 3040 from ASTER and 36 out of 2766 from DAISY in BRCA; 90 out of 1278 from ASTER and 5 out of 1228 from DAISY in STCA.

**Table 4:** Number of Partner genes with Loss-of-Function (LoF) feature in GDSC and proportion associated with drug sensitivity for BRCA (above) and STCA (below).

|  |  | Predicted SL | LoF | Sensitive | Not Sensitive |
| --- | --- | --- | --- | --- | --- |
| BRCA | ASTER | 3040 | 47 | 37 (78.8%) | 10 (21.2%) |
|  | DAISY | 2766 | 36 | 26 (72.2%) | 10 (27.8%) |
|  | ISLE | 0 | - | - | - |
| STCA | ASTER | 1278 | 90 | 90 (100%) | 0 (0%) |
|  | DAISY | 1228 | 5 | 5 (100%) | 0 (0%) |
|  | ISLE | 0 | - | - | - |

Among the pairs with LoF partner genes, ASTER finds more target genes (nearly 79%) that are associated with drug sensitivity compared to DAISY (about 72%), for the test on BRCA cell lines. This shows that among the predicted pairs with LoF partner genes from ASTER, more cell lines showed negative cancer growth when the target genes were targeted by a suitable drug, compared to the predicted cases from DAISY. In the case of STCA cell lines, the number of LoF partner genes among the predicted SL pairs is much lesser in DAISY (5) compared to 90 from ASTER. However, for these LoF partner genes in the predicted pairs, both the methods have 100% association with drug sensitivity. These results suggest that the mutual exclusivity pattern directly mined in ASTER can find SL pairs with potential clinical relevance. However, larger datasets of drug response screens are required for a conclusive evaluation.

#### 4.1.4 Drug Efficacy Prediction

We consider the task of drug efficacy prediction following the experimental procedure in [27, 28]. The principle behind the experiment is that in a SL pair (A,B), loss of function in A leads to *dependence* on B for cell survival. Thus, a drug targeting B would be more effective in cell lines where there is a mutation or copy number alteration causing loss of function in A, compared to those cell lines where there is no loss of function in A. If we measure loss of function by gene expression or SCNA level, then we expect to see a correlation with respect to drug efficacy across cell lines, i.e., the lower the expression and/or SCNA value in A, the higher the drug efficacy of a drug targeting B. Thus, the number of drugs which show such correlations is an indirect way to validate the predicted SL interactions – better SL detection is expected to yield higher number of such drugs.

Similar to the previous experiment, we consider 26 target genes in the in-vitro drug response screen of 24 drugs on 500 cancer cell lines in CCLE. For each method (ASTER, DAISY and ISLE) we (separately) consider the top 5 and 10 predicted SL partner genes, based on the most significant p-values. We again do not consider ASTER++ as this experiment is also designed to evaluate the mutual exclusivity pattern tested by ASTER. We consider two cases to measure loss-of-function (LoF): (i) where the partner gene has low gene expression and (ii) where the partner gene has both low gene expression and low SCNA value. In each case we consider those cell lines with less than median level of expression or normalized SCNA level.

Let *I* be a 500-dimensional vector where each coordinate indicates the number of partner genes (among top 5 and 10, separately), that have been selected for loss-of-function based on the measures mentioned above. Let *J* be the drug efficacy values for a drug that is known to inhibit the target gene in the 500 cell lines. Drug efficacy is given in GDSC in IC50 values which indicates the drug concentration at which a cell line exhibits an absolute inhibition in growth of 50%, lower IC50 implies higher drug sensitivity. The two-sided Spearman correlation p-value between *I* and *J* is then used to measure the correlation between *I* and *J*.

A drug is selected if the correlation test yields a p-value of 0.05 or lesser. We compare the number of drugs selected for each method. Thus, each selected drug represents those where high efficacy is observed on those cell lines containing the drug’s target gene and loss of function of the partner gene. In addition, instead of using a single p-value cutoff of 5%, we use the Benjamini-Hochberg false discovery rate (FDR) controlling procedure [6] with varying FDR thresholds (10%, 20%, 30%, 40% and 50%) to find the number of unique drugs found through the correlations described above.

#### Results

Table 5 shows the number of drugs where high efficacy is observed on those cell lines containing the drug’s target gene and loss of function of the partner gene for SL pairs predicted by ASTER and DAISY. No candidate SL pair passes the tests of ISLE. We find that both ASTER and DAISY yield comparable results. However, when a fixed p-value cutoff is not used, we observe better discrimination between the methods. Fig. 6 shows the number of drugs at varying FDR thresholds using the Benjamini-Hochberg procedure for the same p-values obtained from our correlation test. We find that for Breast Cancer, ASTER outperforms all the baselines, at all 5 FDR values, in terms of the number of drugs. In the case of Stomach Cancer, ASTER has the best performance when only gene expression is considered for the experiment and DAISY has the best performance when both SCNA and gene expression are considered. These results show that the mutual exclusivity pattern mined by ASTER provides a robust signal that is likely to indicate clinically actionable SL pairs.

**Table 5:**
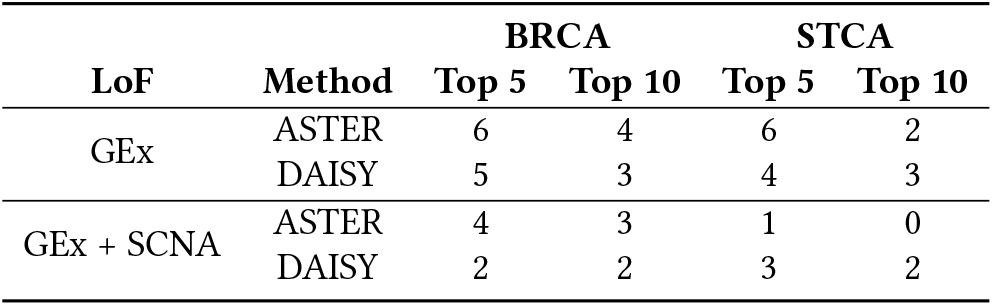
Number of drugs where loss-of-function (LoF) in Top 5 and Top 10 predicted SL partners is correlated (using p-value cutoff 0.05) with drug efficacy with respect to the target gene of the SL pair. LoF is measured by low gene expression (GEx) or both low GEx and low SCNA level (GEx + SCNA). Left: BRCA; Right: STCA.

| LoF | Method | BRCA |  | STCA |  |
| --- | --- | --- | --- | --- | --- |
|  |  | Top 5 | Top 10 | Top 5 | Top 10 |
| GEx | ASTER | 6 | 4 | 6 | 2 |
|  | DAISY | 5 | 3 | 4 | 3 |
| GEx + SCNA | ASTER | 4 | 3 | 1 | 0 |
|  | DAISY | 2 | 2 | 3 | 2 |

**Figure 6:**
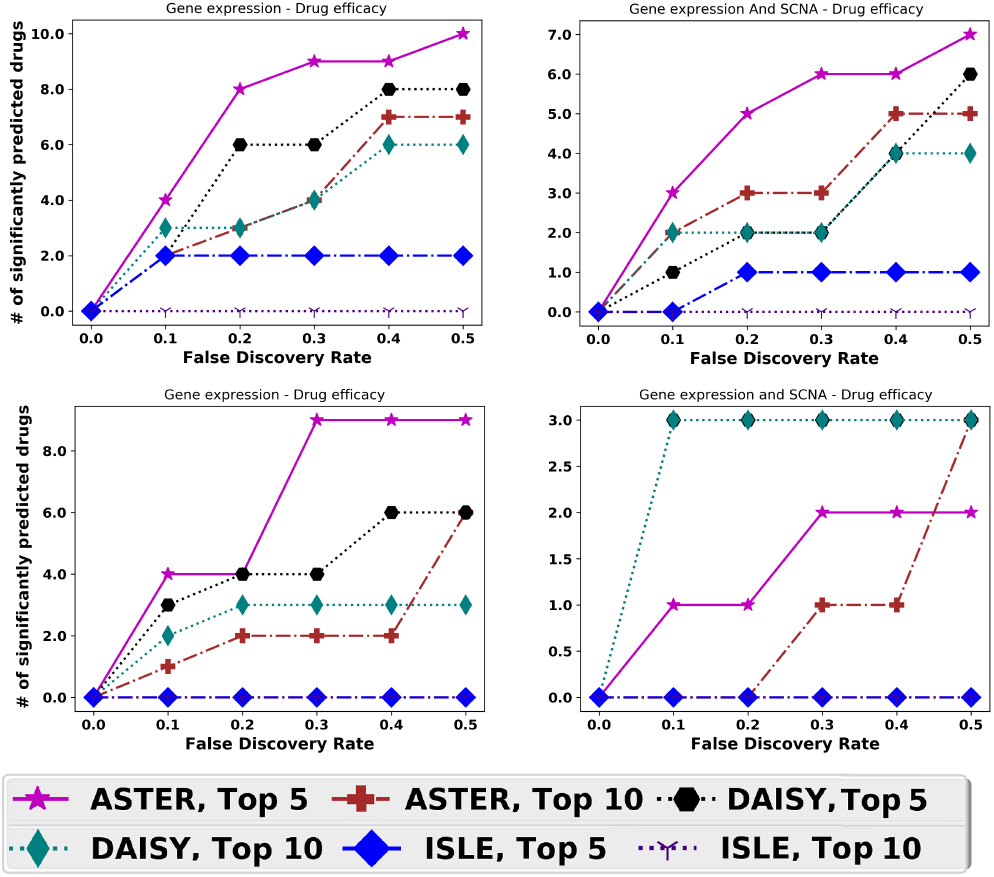
Number of drugs among the 24 drugs with 500 human cancer cell lines, whose efficacy is predicted with statistical significance at varying FDR levels based on correlation of drug efficacy on target gene with loss of function [measured using gene expression alone (left) and with SCNA (right)] of top 5/10 predicted partner genes from ASTER, DAISY and ISLE. Top: BRCA; Bottom: STCA.

### 4.2 Qualitative Analysis of Identified SL pairs

We analyze the predicted pairs from ASTER in breast and stomach cancer. We use the same 16,916 gene pairs listed in SynLethDB [21] as input. However, different tissue-specific samples are used from TCGA and GTEx for each cancer. A p-value threshold of 0.01 is used for each test in ASTER. Tables 6 and 7 list the top pairs of genes, in Breast and Stomach cancer respectively, selected by ASTER. Notably, ASTER was able to identify the well-known BRCA-PARP pair that DAISY and ISLE, in their default settings with this input data, could not find.

**Table 6:** Breast Cancer gene pairs selected by ASTER from candidates in SynLethDB with all 3 p-values *<* 0.01.

| Gene A |  | Gene B |  | P1 | P2 | P3 |
| --- | --- | --- | --- | --- | --- | --- |
| PARP1 | ↑ | BRCA2 | ↓ | 2.61E-39 | 1.33e-04 | 1.88e-04 |
| BRCA2 | ↓ | PARP1 | ↑ | 5.86E-08 | 1.33e-04 | 1.19e-03 |
| DHX9 | ↑ | ESCO2 | ↓ | 1.23E-34 | 6.17e-04 | 8.32e-04 |
| ESCO2 | ↓ | DHX9 | ↑ | 9.21E-22 | 6.17e-04 | 1.17e-03 |
| MED4 | ↓ | SRP9 | ↑ | 1.29E-06 | 6.17e-04 | 2.94e-03 |
| TNFRSF10B | ↓ | FADD | ↑ | 1.16e-03 | 6.84E-06 | 4.50e-04 |
| ILF2 | ↑ | RFC3 | ↓ | 2.59E-40 | 3.04e-03 | 3.39e-03 |
| S100A2 | ↑ | GPX8 | ↓ | 8.04E-19 | 2.00e-03 | 6.15e-03 |
| CAPN2 | ↑ | GPX8 | ↓ | 2.92E-15 | 5.34e-03 | 3.41e-03 |
| IKBKB | ↑ | TNFRSF10A | ↓ | 1.66E-10 | 1.41e-03 | 3.58e-04 |
| PTPN14 | ↑ | CTSB | ↓ | 4.54e-04 | 3.54e-03 | 3.48e-03 |

**Table 7:** Stomach Cancer gene pairs selected by ASTER from candidates in SynLethDB with all 3 p-values *<* 0.01.

| Gene A |  | Gene B |  | P-Value1 | P-Value2 | P-Value3 |
| --- | --- | --- | --- | --- | --- | --- |
| ABCC10 | ↑ | CDKN2A | ↓ | 1.12e-14 | 3.19e-04 | 2.01e-03 |
| CDKN2A | ↓ | ABCC10 | ↑ | 4.60e-13 | 1.36e-04 | 2.56e-04 |
| CDKN2A | ↓ | SLC29A1 | ↑ | 4.60e-13 | 2.21e-03 | 8.09e-05 |
| SLC29A1 | ↑ | CDKN2A | ↓ | 1.01e-09 | 1.89e-04 | 1.89e-04 |
| KRAS | ↑ | MACROD2 | ↓ | 4.38e-16 | 4.12e-03 | 5.05e-04 |
| MACROD2 | ↓ | KRAS | ↑ | 3.00e-13 | 1.36e-04 | 3.27e-04 |
| MYC | ↑ | RAD51B | ↓ | 5.04e-17 | 3.76e-03 | 4.11e-03 |

SL pairs tend to participate in closely related biological processes [16, 30, 48]. This also forms the basis of the third test in DAISY. Functional correlation between genes in a predicted SL pair provides an indirect indicator of SL. To test this, we obtain GO terms associated with each gene in the predicted pairs (in Tables 6 and 7), using DAVID version 6.8 [25] with respect to Gene Ontology (GO) — Biological Processes (BP). For a pair of genes (A,B), we denote the sets of functional annotation terms found for A and B by *F*_*A*_ and *F*_*B*_ respectively. To reduce the effects of potential annotation errors [3, 8, 19], we consider only those pairs where each gene A, B has at least 50 annotation terms. We compute the union *F*_*A*_ ∪ *F*_*B*_ and intersection *F*_*A*_ ∩ *F*_*B*_ of these sets and find the Jaccard Index given by |*F*_*A*_ ∩ *F*_*B*_ |/|*F*_*A*_ ∪ *F*_*B*_ | where |.| denotes set cardinality.

Table 8 shows the number of annotation terms for each gene, the union and intersection for each pair from which the Jaccard indices are calculated. We find that most pairs have indices greater than 0.2 that is similar to that of PARP/BRCA pair. Tables 10 and 11 in the Appendix show functional annotations, with respect to KEGG Pathways of the genes in the most significant predicted SL pairs – we consider 90 genes for Breast Cancer and 24 genes for Stomach Cancer (with p-value *<* 0.05). We observe that the associated pathways are known to be related to cancer. These results show that through the mutual exclusivity pattern mined by ASTER, signals of functional correlations are found, although ASTER does not explicitly search for it.

**Table 8:** Jaccard Index (𝔍) of GO — BP functional annotation terms for genes in predicted SL pairs (from Tables 6 and 7).

| Gene A | Gene B | $\mathcal{J}$ | $\vert F_A \cap F_B\vert $ | $\vert F_A \cup F_B\vert $ | $\vert F_A\vert $ | $\vert F_B\vert $ |
| --- | --- | --- | --- | --- | --- | --- |
| PARP1 | BRCA2 | 0.238 | 149 | 624 | 492 | 288 |
| DHX9 | ESCO2 | 0.266 | 32 | 120 | 72 | 80 |
| MED4 | SRP9 | 0.193 | 32 | 166 | 98 | 102 |
| TNFRSF10B | FADD | 0.271 | 126 | 465 | 169 | 428 |
| ILF2 | RFC3 | 0.357 | 49 | 137 | 69 | 120 |
| TNFRSF10A | IKBKB | 0.267 | 96 | 360 | 295 | 163 |
| PTPN14 | CTSB | 0.102 | 21 | 205 | 108 | 119 |
| MYC | RAD51B | 0.097 | 75 | 776 | 730 | 124 |

**Table 9:** TP, FP, TN and FN counts within each test and final results on the Leukamia dataset Pathway Name.

|  | DAISY |  |  |  | ISLE |  |  |  |
| --- | --- | --- | --- | --- | --- | --- | --- | --- |
|  | TP | FP | TN | FN | TP | FP | TN | FN |
| 1st | 130 | 24 | 15289 | 15183 | 94 | 50 | 15263 | 15219 |
| 2nd | 1654 | 619 | 14694 | 13659 | 0 | 0 | 15313 | 15313 |
| 3rd | 2526 | 831 | 14482 | 12787 | 5740 | 9492 | 5821 | 9573 |
| final | 4 | 0 | 15313 | 15309 | 0 | 0 | 15313 | 15313 |

|  | ASTER |  |  |  | ASTER++ |  |  |  |
| --- | --- | --- | --- | --- | --- | --- | --- | --- |
|  | TP | FP | TN | FN | TP | FP | TN | FN |
| 1st | 654 | 629 | 14684 | 14659 | 654 | 629 | 14684 | 14659 |
| 2nd | 0 | 0 | 15313 | 15313 | 0 | 0 | 15313 | 15313 |
| 3rd | 0 | 0 | 15313 | 15313 | 0 | 0 | 15313 | 15313 |
| final | 0 | 0 | 15313 | 15313 | 0 | 0 | 15313 | 15313 |

**Table 10:** DAVID 6.8 Pathway Enrichment Analysis (KEGG) for predicted Top 90 SL pairs (BRCA) from SynLethDB by ASTER with p-value *<* 0.05.

| Pathway Name | # Genes | % | P-value | Benjamini |
| --- | --- | --- | --- | --- |
| Pathways in cancer | 19 | 19.4 | 5.8E-10 | 9.3E-8 |
| Apoptosis | 9 | 9.2 | 2.4E-8 | 1.9E-6 |
| p53 signaling pathway | 6 | 6.1 | 1.8E-4 | 2.0E-3 |
| PI3K-Akt signaling pathway | 10 | 10.2 | 1.5E-3 | 1.1E-2 |
| Cell cycle | 6 | 6.1 | 3.0E-3 | 2.0E-2 |
| MAPK signaling pathway | 8 | 8.2 | 3.8E-3 | 2.4E-2 |
| ErbB signaling pathway | 5 | 5.1 | 5.0E-3 | 3.0E-2 |
| Thyroid hormone signaling pathway | 5 | 5.1 | 1.3E-2 | 6.8E-2 |
| Neurotrophin signaling pathway | 5 | 5.1 | 1.5E-2 | 7.5E-2 |
| Natural killer cell mediated cytotoxicity | 5 | 5.1 | 1.6E-2 | 7.7E-2 |
| Homologous recombination | 3 | 3.1 | 2.2E-2 | 1.0E-1 |
| Ras signaling pathway | 6 | 6.1 | 3.4E-2 | 1.4E-1 |
| T cell receptor signaling pathway | 4 | 4.1 | 4.5E-2 | 1.8E-1 |

**Table 11:** DAVID 6.8 Pathway Enrichment Analysis (KEGG) for predicted Top 24 SL pairs (STAD) from SynLethDB by ASTER with p-value *<* 0.05.

| Pathway Name | # Genes | % | P-value | Benjamini |
| --- | --- | --- | --- | --- |
| Pathways in cancer | 8 | 30.8 | 1.4E-5 | 7.6E-4 |
| Homologous recombination | 3 | 11.5 | 2.0E-3 | 2.0E-2 |
| Foxo signaling pathway | 4 | 15.4 | 3.4E-3 | 3.0E-2 |
| MicroRNAs in cancer | 5 | 19.2 | 3.6E-3 | 3.0E-2 |
| Signaling pathways regulating pluripotency of stem cells | 4 | 15.4 | 3.8E-3 | 3.0E-2 |
| p53 signaling pathway | 3 | 11.5 | 1.0E-2 | 6.1E-2 |
| ErbB signaling pathway | 3 | 11.5 | 1.7E-2 | 8.5E-2 |
| Cell cycle | 3 | 11.5 | 3.3E-2 | 1.3E-1 |

In addition to BRCA and PARP, genes in the other identified breast cancer SL pairs, through the most significant p-values in ASTER, are also known to play important roles in cancer. DHX9 is known to block the DNA repair function of BRCA-1, leading to cancer growth [13]. ESCO2 is cell cycle-related gene involved in cancer progression [56]. FADD is a common mediator of apoptosis. TRNFRSF10B and TRNFRSF10A induce FADD-dependent apoptosis [14]. Both TRNFRSF10A and TRNFRSF10B are potent stimulators of apoptosis and promising targets for cancer therapy [29]. IKBKB gene is known to prevent apoptosis in some contexts [44]. In stomach cancer, CDKN2A is tumor suppressor gene and hypermethylation of CDKN2A in gastric cancer correlates to enhanced tumorigenesis and poor prognosis [59]. KRAS is a well-known oncogene that regulates growth and proliferation. MACROD2 deletion in human colorectal cancer was shown to cause impaired PARP1 activity and chromosome instability [41]. MYC is proto-oncogene that regulates cell cycle, differentiation, proliferation, and is highly amplified in tumor samples. MYC is known to modulate cell survival in response to DNA-damaging agents and is a potential target gene [36].

These results suggest that through the mutual exclusivity pattern mined, ASTER is able to identify pairs with potential therapeutic value. More investigations are required for a deeper understanding of how these patterns are related to the underlying biological contexts in which these genes promote or disrupt cancer progression.

## 5 CONCLUSION

In this paper, we presented ASTER, a technique based on hypothesis testing, to identify SL pairs that leverages unified tissue-specific gene expression data from GTEx and TCGA. We also discussed how it can be extended, through the use of AdaFDR [58], for largescale multiple hypothesis testing and to adaptively find a decision threshold based on additional input gene features. ASTER identifies SL in an input gene pair through the application of 3 simple tests using only RNA-Seq and SCNA data. These tests are designed to mine the mutual exclusivity pattern that can potentially identify SL. ASTER++ enables the use of additional gene features, when available, within the interpretable hypothesis testing framework.

We evaluated ASTER through multiple experiments. ASTER and ASTER++ outperformed state-of-the-art methods based on hypothesis testing (DAISY and ISLE) in identifying true SL pairs on two benchmark datasets. We evaluated the predicted pairs through their clinical actionability that is implied by the property of Synthetic Lethality. The number of clinically actionable pairs identified by ASTER were higher than those identified by DAISY and ISLE. We also qualitatively analyzed the genes from predicted pairs from ASTER with respect to their functional annotations and roles in cancer. These experiments suggest that the mutual exclusivity pattern mined by ASTER provides a strong signal to identify SL pairs. An implementation of ASTER is available at https://github.com/lianyh/ASTER.

In the future, we plan to validate our predictions for previously untested gene pairs through CRISPR screens. Future work can also further explore ways to incorporate additional drug-related information to improve the clinical actionability of ASTER.

## APPENDIX

## Notes

### Competing Interest Statement

The authors have declared no competing interest.

